# Water mass age structures the auxiliary metabolic gene content of free-living and particle-attached deep ocean viral communities

**DOI:** 10.1101/2022.10.13.512062

**Authors:** Felipe H Coutinho, Cynthia B Silveira, Marta Sebastián, Pablo Sánchez, Carlos M Duarte, Dolors Vaqué, Josep M Gasol, Silvia G Acinas

## Abstract

Viruses play important roles on the biogeochemical cycles that take place in the ocean.Yet, deep ocean viruses are one of the most under-explored fractions of the global biosphere. Little is known about the environmental factors that control the composition and functioning of their communities, or how they interact with their free-living or particle-attached microbial hosts. Thus, we analysed 58 viral communities associated to size fractionated free-living (0.2–0.8 μm) and particle-attached (0.8–20 μm) cellular metagenomes from bathypelagic (2,150-4,018 m deep) microbiomes obtained during the Malaspina expedition. These metagenomes yielded 6,631 viral sequences, 91% of which were novel, and 67 represented high-quality genomes. Taxonomic classification assigned 53% of the viral sequences to families of tailed viruses from the order Caudovirales. Computational host prediction associated 886 viral sequences to dominant members of the deep ocean microbiome, such as Alphaproteobacteria (284), Gammaproteobacteria (241), SAR324 (23), Marinisomatota (39), and Chloroflexota (61). Free-living and particle-attached viral communities had markedly distinct taxonomic composition, host prevalence, and auxiliary metabolic gene content, which led to the discovery of novel viral encoded metabolic genes involved in the folate and nucleotide metabolisms. Water mass age emerged as an important factor driving viral community composition. We postulated this was due to changes in quality and concentration of dissolved organic matter acting on the host communities, leading to an increase of viral auxiliary metabolic genes associated with energy metabolism among older water masses. These results shed light on the mechanisms by which environmental gradients of deep ocean ecosystems structure the composition and functioning of free-living and particle-attached viral communities.

## Introduction

Aquatic viruses play major roles in the structuring of microbiomes and modulate biogeochemical cycles of global relevance [1,2]. Global scale studies based on viral metagenomics have brought significant advances to our understanding of viral genomic diversity and the environmental factors that structure their communities [3–5]. These studies most often evaluated viruses from the epipelagic and mesopelagic zones, while the bathypelagic received less attention, although some notable exceptions led to significant advances in the field [6–9]. Nevertheless, our understanding of viral diversity in the largest marine ecosystem, the deep ocean, is still limited [6,7,10]. Compared to the epipelagic zone, the bathypelagic is characterized by the absence of light, low temperatures, very low concentrations of labile carbon, and higher concentrations of inorganic nutrients [11]. Also, the bathypelagic has lower densities of prokaryotic cells and viral particles but higher virus-to-prokaryote ratio [11–13]. In the bathypelagic both free-living and particle-attached microbial communities are active, but differ regarding their taxonomic and functional composition, cell densities, and activity levels [14,15]. These differences affect the structure and functioning of the viral communities that infect them [8]. Furthermore, evidence suggests that deep ocean viruses contribute to organic matter remineralization through lysis of particle-attached heterotrophic hosts, possibly enhancing carbon export efficiency [16]. Yet, there is little information regarding which are the hosts targeted by these viruses, their genetic diversity, and the mechanisms by which they interact with their host communities.

Viral genomes often encode auxiliary metabolic genes (AMGs) that re-direct host metabolism during infection towards pathways that benefit viral replication [4,17]. Changes in host metabolism brought by the expression of AMGs impact global element and energy cycles [12,18]. The deep ocean has a unique set of AMGs [19–21]. Yet the processes driving the composition of this genetic repertoire have not been assessed with emphasis on differences in physical and chemical parameters that characterize different water masses of the deep oceans. Likewise, the differences in viral taxonomy, host prevalence, and AMG content between viruses associated with the free-living and particle-attached communities have not been explored in detail.

The Malaspina expedition has contributed to the effort to describe microbial diversity and functioning in the oceans [14,22]. Findings derived from this expedition revealed that deep ocean basins and water masses have a major role in structuring the composition of bathypelagic communities of bacteria, archaea, and micro-eukaryotes [22,23]. Bathypelagic free-living (FL) and particle-attached (PA) communities have markedly distinct taxonomic [24], and functional compositions [14]. While FL microbial assemblages are more diverse and contain more oligotrophic taxa, the PA microbial assemblages are less diverse and contain more copiotrophic taxa [24], although some taxa occur in both fractions, displaying a dual lifestyle [15]. Similarly, viral community diversity is different across viral and cellular size fractions [6,25]. Most studies of bathypelagic viruses have focused on free viral particles in the smallest size fraction (< 0.22 μm) [10,26,27]. Less is known regarding the environmental parameters that control the structure and functioning of the viral communities in the cellular fraction (> 0.22 μm), i.e. those associated to particle-attached or free-living host communities. Based on previous findings, we hypothesize that bathypelagic free-living and particle-attached viral communities differ as a consequence of the differences in host community composition and functioning. We postulate that particle-attached communities are enriched in viruses that target copiotrophic bacteria, as particles are considered resource rich micro-environments within the bathypelagic, where labile organic matter sources predominantly are depleted. We further hypothesize that the environmental parameters that characterize different water masses in the deep ocean also shape the taxonomic composition and AMG content of bathypelagic viral communities, specifically regarding metabolic pathways associated with energy and resource availability.

## Results and Discussion

### Novel viruses from the bathypelagic zone have a unique genetic repertoire

The Malaspina samples were obtained to represent the tropical and subtropical bathypelagic ocean (Figure 1A and Table S1). Three main water masses were identified, which differed according to salinity, temperature, and Apparent Oxygen Utilization (AOU): Circumpolar Deep Water (CDW), North Atlantic Deep Water (NADW), and Weddell Sea Deep Water (WSDW) [28], (Figure 1B). The AOU variable integrates all respiratory processes since water mass formation [29], thus older water masses display higher values of apparent oxygen utilization. Cellular metagenomes were generated from 28 samples from the free-living (0.2–0.8 μm) fraction and 30 samples from the particle-attached (0.8–20 μm) fraction, for a total of 58 metagenomes (Table S1). Out of 422,928 scaffolds derived from the co-assembly of the metagenomes, VIBRANT [30] classified 6,631 scaffolds as fully viral (6,479) or as viral fragments (152) within longer scaffolds (Table S2). Among these, CheckV [31] categorized 23 as complete genomes and 44 as high-quality genome fragments (i.e., estimated completeness ≥ 90%). VPF-Class [32] classified 5,100 scaffolds as dsDNA viruses and 33 as ssDNA viruses. The most common families were Myoviridae (1,856), Siphoviridae (1,039), Podoviridae (637) and Phycodnaviridae (86). The most common taxa assigned as putative hosts of the viral scaffolds by PHIST [33] were Alphaproteobacteria (284), Gammaproteobacteria (241), Chloroflexota (61), Bacteroidota (60), and Marinisomatota (39).

**Figure 1:**
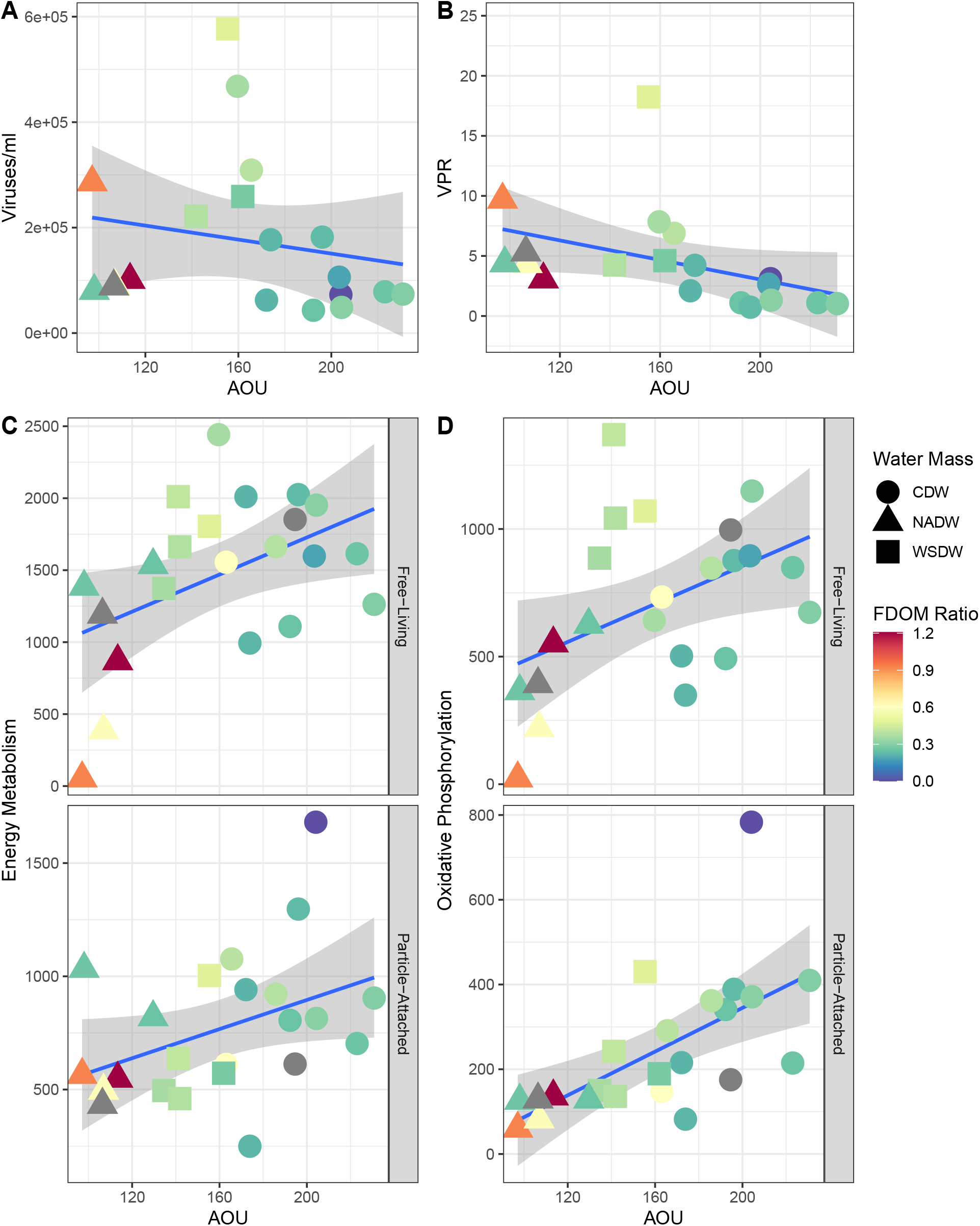
Global trends of the deep ocean viral communities. A) World map displaying sampling sites from which Malaspina bathypelagic metagenomes were retrieved. B) Scatter plot displaying the values of salinity, temperature, and AOU measured at the sites from which metagenome samples were retrieved. C) NMDS displaying the separation of samples by size fraction (NMDS1) and AOU (NMDS2). Sample MP0262 was excluded from this analysis as it displayed extremely high RPKM values, not in line with those observed in any other samples.

The 563,348 Coding DNA Sequences (CDS) derived from the viral genomes and genome fragments were annotated against the UniRef. Pfam, and KEGG databases (Table S3). Also, all CDS were queried against a custom reference database of isolated and uncultured viral genomes encompassing multiple ecosystems [34]. Pairwise similarity was calculated between Malaspina viral scaffolds and reference database genomes, based on the percentage of matched CDS. Only 598 (9%) of the Malaspina viral scaffolds shared 10 or more CDS with a single sequence in the reference database. These results suggest that 91% of Malaspina viral sequences are novel, while the remaining ones only have distant relatives in reference databases.

### Free-living and particle-attached metagenomes have unique viral communities

Viral scaffold relative abundances (Reads Per Kilobase per Megabase of metagenome, RPKM) were calculated by read mapping to determine the viral community composition in the metagenomes (Table S4). Free-living samples and particle-attached viral communities were distinguished by the first axis of a non-metric multidimensional scaling analysis (Figure 1C, NMDS stress = 0.11). Differences in viral community composition between FL and PA samples were significant according to PERMANOVA (*p* < 0.001). The second NMDS axis separated samples between lower and higher Apparent oxygen utilization (AOU). The AOU of the three water masses differed according to their age: NADW (median age = 481 years, median AOU = 107 μmol kg^-1^), WSDW (median age = 545 years, median AOU = 142 μmol kg^-1^), and CDW (median age = 1,046 years, median AOU = 193 μmol kg^-1^). Apparent oxygen utilization is an important factor associated with host community composition in the bathypelagic [15,22]. Our results corroborate these findings and extend them by showing that water mass age is a significant driver of bathypelagic viral community composition.

We next analysed how viral community composition shifted between free-living and particle-attached fractions. We calculated the relative abundances of viruses grouped by taxonomic family (Table S8). Families Myoviridae, Siphoviridae, and Podoviridae were the dominant taxa across all samples, regardless of fraction or water mass (Figure 2A). Yet, the families Myoviridae and Siphoviridae were significantly more abundant among FL samples (Mann-Whitney test, *p* < 0.01), while families Podoviridae and Phycodnaviridae were significantly more abundant among PA samples (Mann-Whitney test, *p* < 0.01). Phycodnaviridae only had abundances above 2,500 RPKM in the particle-attached fraction (except for station 62). Viruses of the family Phycodnaviridae infect eukaryotic microalgae, hence their higher abundance among samples from the larger size fraction, as sinking particles may be composed by phytodetritus [35], or even intact phytoplankton cells [36].

**Figure 2:**
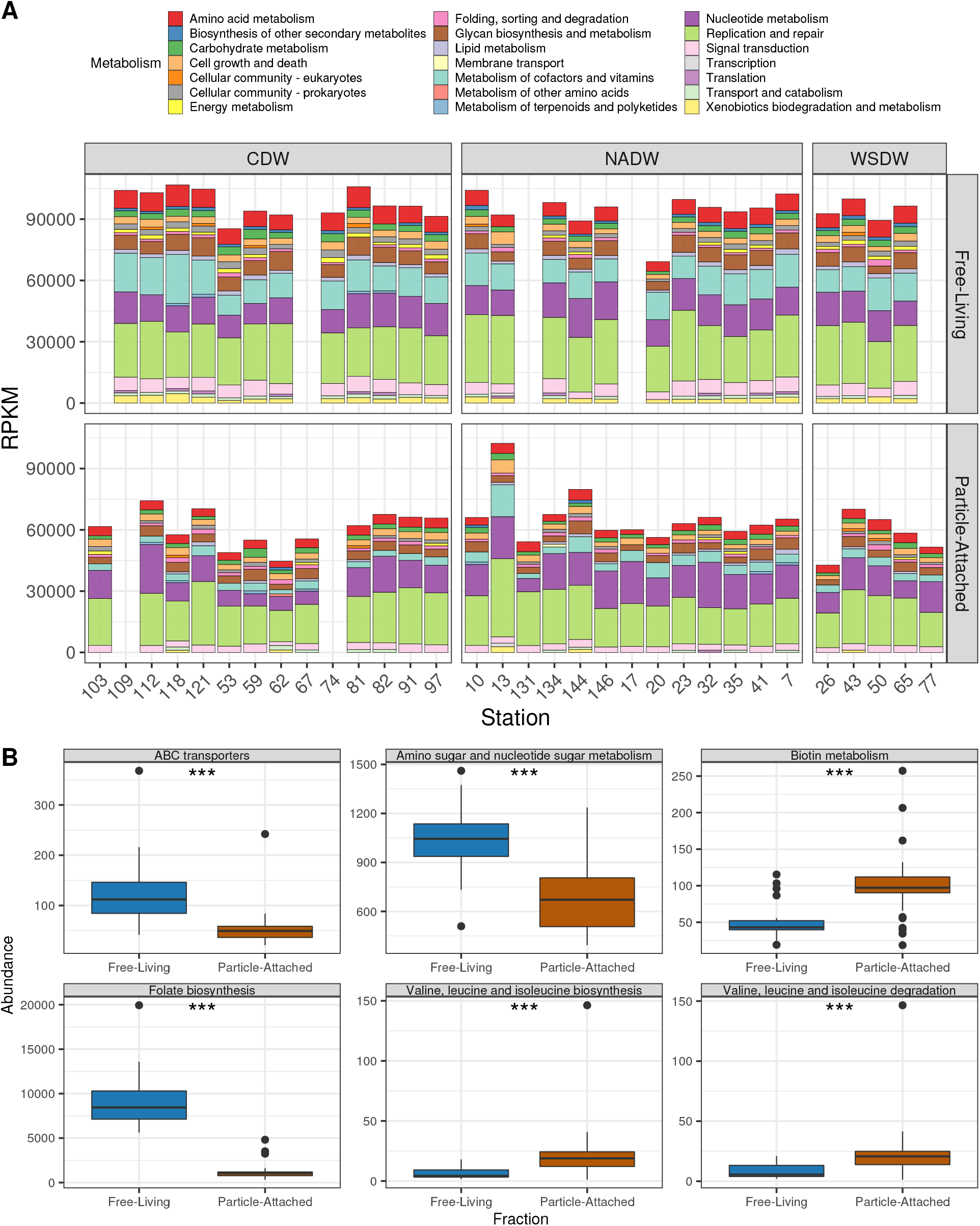
Differences in viral community taxonomic composition between free-living and particle-attached samples. A) RPKM abundances of viral genomes grouped by family level taxonomic classification. B) RPKM abundances of viral genomes grouped by predicted host phylum (or class for Proteobacteria). Sampling stations are sorted from left to right by increasing oxygen concentrations. Sample MP0262 was excluded from this analysis as it displayed extremely high RPKM values, not in line with those observed in any other samples.

We calculated the relative abundances of viruses grouped by predicted host phylum (or class for Proteobacteria, Table S9). Viruses predicted to infect Alphaproteobacteria and Gammaproteobacteria were among the most abundant ones in both PA and FL samples (Figure 2B). Nevertheless, multiple viral groups displayed significantly different relative abundances among PA and FL samples (Mann-Whitney test, *p* < 0.01). Viruses predicted to infect Marinisomatota, Chloroflexota, and SAR324 were more abundant among FL samples, while viruses predicted to infect Myxococcota, Planctomycetota, Actinobacteriota were more abundant among PA samples. These trends, are in agreement with the previously observed taxonomic composition of the host communities reported for the same samples [14,15,22,24]. We postulate that differences in target host prevalence between the two fractions are a consequence of differences in the preferred ecological niche of deep ocean microbes, which can be roughly divided into particle-attached copiotrophs (e.g. Gammaproteobacteria and Actinobacteriota), free-living oligotrophs (e.g. SAR324 and Chloroflexota), and those that present a dual lifestyle [14].

### Viral auxiliary metabolic gene content shifts between free-living and particle-attached fractions

All metabolic pathways of auxiliary metabolic genes were relatively more abundant in the FL than the PA fraction (Figure 3A and Table S7). This trend was most evident for the metabolism of cofactors and vitamins, which was the main AMG category responsible for the differences in relative abundances between the fractions. Out of the 309 AMGs involved in the metabolism of cofactors and vitamins, only 54 were derived from viral scaffolds with a predicted host, which most often were Alphaproteobacteria (20) and Gammaproteobacteria (14). In addition, AMGs from the carbohydrate, amino acid, lipid, and energy metabolisms were also more abundant in the FL than the PA samples. We propose that in PA communities, where more labile organic matter is available, viruses have less necessity to manipulate host metabolism during infection. Meanwhile, in the FL communities, which have less access to labile organic matter, viruses would more often encode AMGs, manipulating host metabolism towards successful production of viral progeny during lytic infection.

**Figure 3:**
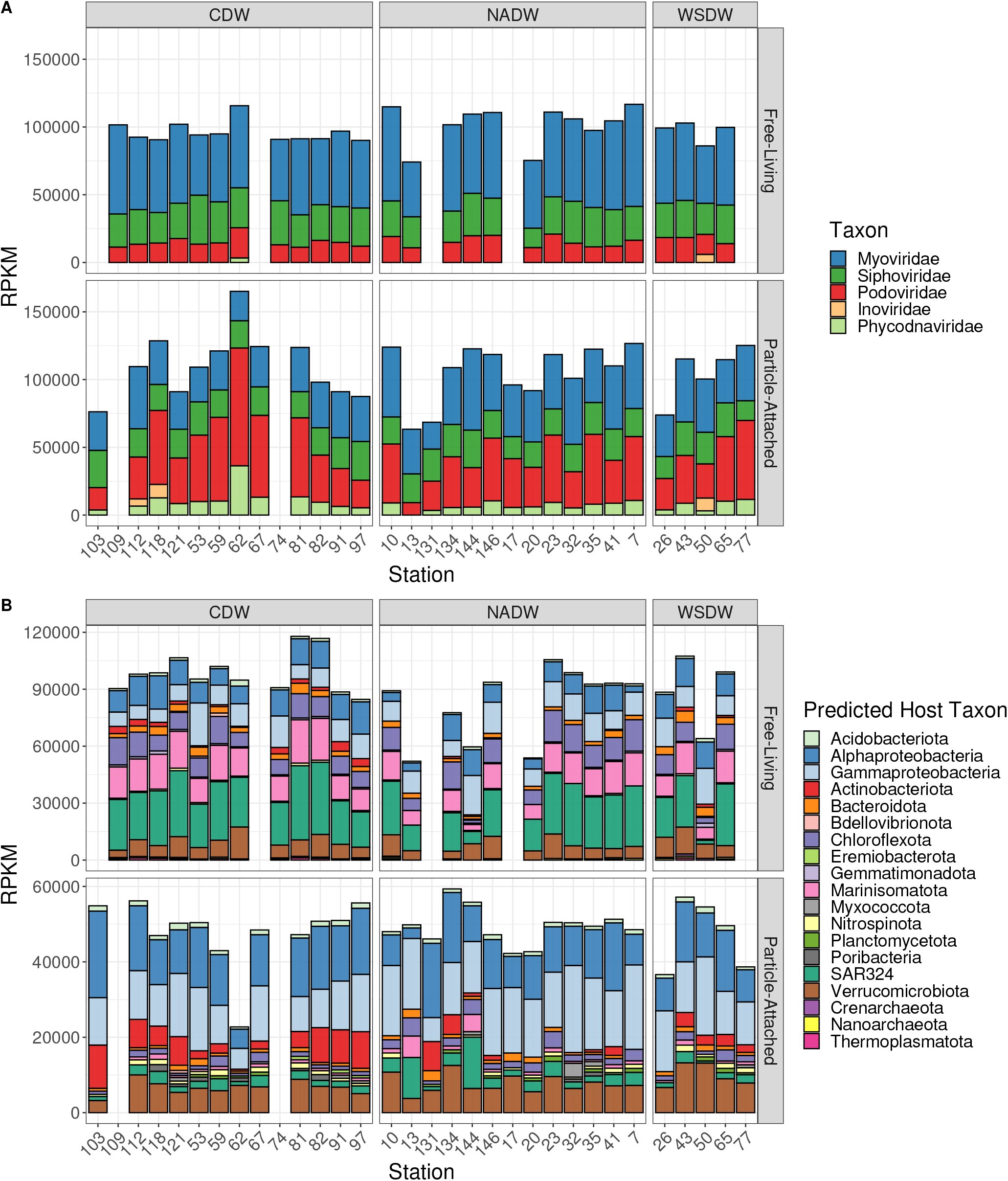
Differences in viral community functional composition between free-living and particle-attached samples. A) Auxiliary metabolic gene KEGG module abundances across samples Sampling stations are sorted from left to right by increasing oxygen concentrations. B) Boxplots depicting the differences in pathway abundances between free-living and particle-attached fractions. Boxes depict the median, the first and third quartiles. Whiskers extend to 1.5 of the interquartile ranges. Outliers are represented as dots above or below whiskers. The triple asterisks indicate *p*-values< 0.001 obtained with the Mann-Whitney test. Sample MP0262 was excluded from this analysis as it displayed extremely high RPKM values, not in line with those observed in any other samples.

This finding corroborates the observation that vitamin biosynthesis genes are enriched in FL microbial metagenomes compared to PA metagenomes obtained from the same Malaspina samples [15], and indicates that this pattern also extends to the viral fraction. Regarding specific metabolic pathways, AMGs involved in ABC transporters; amino sugar and nucleotide sugar metabolism; and folate biosynthesis enzymes were more abundant in FL samples (corrected *p-*value ≤ 0.05, Figure 3B and Table S6). Biotin metabolism and the biosynthesis and degradation pathways of valine, leucine and isoleucine were relatively more abundant among PA samples (corrected *p* ≤ 0.05, Figure 3B).

Given the differential abundance of AMGs involved in the metabolism of cofactors and vitamins, we investigated viral genomes encoding such genes in further detail, to determine the specific mechanisms by which viruses could modulate host vitamin metabolism in the bathypelagic. A total of 14 viral scaffolds had AMGs encoding for a dihydrofolate reductase (DHFR). This enzyme catalyses multiple reactions, including the conversion of folate (Vitamin B9) and 7,8-dihydrofolate to 5,6,7,8-tetrahydrofolate (THF). Only three of the viral scaffolds encoding DHFR could reliably be assigned to a host (*p* ≤ 2.384e-14), one to Gammaproteobacteria and two to Marinisomatota. Among the 14 scaffolds, Malaspina_Vir_6045 was the longest (214 Kbp). This sequence was derived from a viral genome estimated to be 79% complete, and classified as a member of the family Myoviridae. Host prediction would have associated this viral genome to a member of the class Gammaproteobacteria, yet this prediction did not pass our stringent threshold (*p* = 3.6e-04). Aside from DHFR, scaffold Malaspina_Vir_6045 was the only one that encoded multiple other AMGs, which gave us detailed insights about the potential of this virus to interfere with host metabolism (Figure 4A). This viral genome encoded five AMGs involved in dTMP biosynthesis, namely ribonucleoside-diphosphate reductase (subunits alpha and beta), dCTP deaminase, dUTPase, thymidylate synthase, and dTMP kinase. Thymidylate synthase catalyses the conversion of dUMP into dTMP using 5,10-methylene-tetrahydrofolate as a co-substrate, which is subsequently converted to dTDP by dTMP kinase, and finally to dTTP by a nucleoside diphosphate kinase (Figure 4B). The occurrence of these genes in a single genome suggests that this virus has the potential to enhance THF biosynthesis during infection, and re-direct it to the synthesis of deoxynucleotides to be used in viral genome replication.

**Figure 4:**
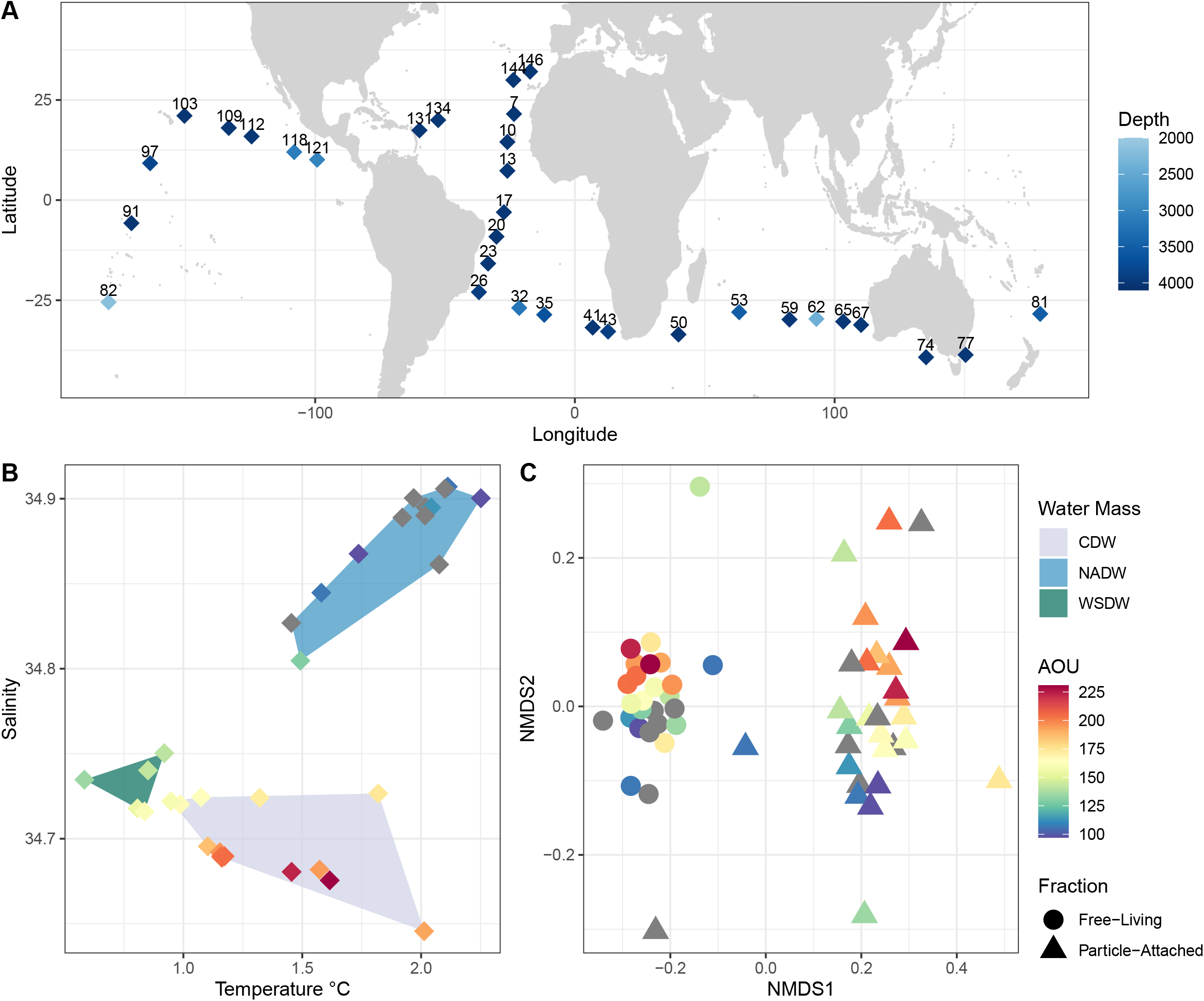
Genomic composition and AMG content of sequence Malaspina_Vir_6045. A) Genomic map depicting the protein-encoding genes (depicted as arrows) identified in the viral scaffold. Genes are colour-coded according to their functional annotation within the five categories: structural genes (red), genome replication genes (green), packaging and lysis genes (yellow), auxiliary metabolic genes (orange), and others (blue). B) Proposed scheme illustrating the metabolic reactions within the folate and nucleotide pathways of the host which are under the influence of viral AMGs encoded in the genomic sequence of Malaspina_Vir_6045. Enzyme names are depicted in red, those genes encoded in the viral genome are enclosed by purple rectangles.

The relative abundances of Malaspina_Vir_6045 ranged between 0 and 223 RPKM, and the mean abundance was 10 RPKM, which falls into the 69^th^ percentile when considering the relative abundances of all scaffolds across all samples. Relative abundances for Malaspina_Vir_6045 were higher among PA samples, specifically those from NADW. When considering DHFR encoding viral scaffolds together, the opposite pattern was observed, as these had a mean abundance of 61 RPKM among FL samples and 23 RPKM among PA samples (Figure S1). The current results provide evidence that the viruses play a role in the budget of vitamins at the bathypelagic. DHFR genes have previously been reported in the genomes of marine bacteriophages [37]. Yet, to our knowledge, this is the first time this gene is reported in bathypelagic viruses, in association with other genes directly involved in nucleotide metabolism, and with differential abundances between free-living and particle-attached fractions or water masses. Thus, it is possible that the abundance of DHFR encoding viruses is partially regulated by the exchange of vitamins between their free-living and particle-attached hosts.

### Viral community abundance, AMG diversity, and composition are associated with water mass age

Viral particle abundances and virus-to-prokaryote ratios, quantified by flow cytometry, were negatively correlated with AOU (Figures 5A and 5B, Spearman correlation coefficient = −0.45, *p* < 0.05 and = −0.72, *p* < 0.001, respectively). Likewise, a significant positive correlation was observed between AOU and the relative abundance of the AMGs related to energy metabolism (Figure 5C, Spearman correlation coefficient = −0.34, *p* < 0.05), and between AOU and AMGs of the oxidative phosphorylation pathway (Figure 5D, Spearman correlation coefficient = −0.43, *p* < 0.005).

**Figure 5:**
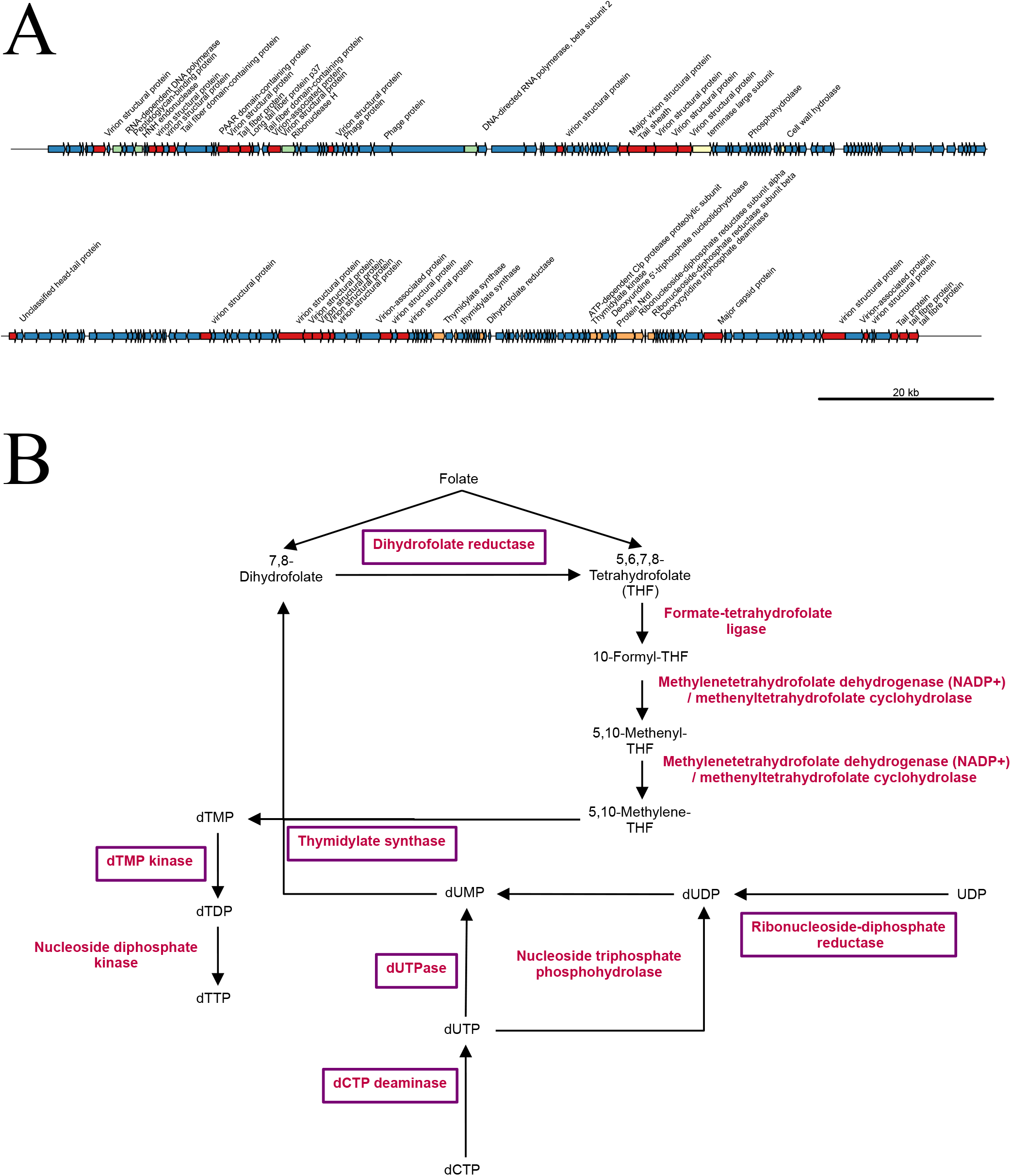
Associations between deep ocean viral communities and AOU. Scatter plots depict the association between AOU (x-axis) and the tested variable (y-axis). The blue line depicts the best fit for linear regression models with shaded areas depicting the standard error. A) Association between viral absolute abundance and AOU. B) Association between virus-to-prokaryote ratio (VPR) and AOU. C) Association between relative abundance of energy metabolism AMGs and AOU in FL and PA samples. D) Association between relative abundance of oxidative phosphorylation AMGs and AOU in FL and PA samples. Sample MP0262 was excluded from this analysis as it displayed extremely high RPKM values, not in line with those observed in any other samples.

The association patterns between AOU and the absolute abundances of viruses, as well as the relative abundances of the AMGs encoded within them, suggest that the composition and functioning of viral communities are influenced by water mass age. Even among the water masses with highest AOU values, it is unlikely that the microbial communities therein are subjected to a limited oxygen supply. Therefore, the observed associations with AOU are likely linked to the differences in organic matter content among water masses. The optical properties of dissolved organic matter can be used as a tracer of biochemical processes [38], distinguishing between humic-like (recalcitrant) versus amino-acid like (labile) fluorescence. Thus, the ratio between labile and recalcitrant fluorescence in a sample (hereafter FDOM ratio) may be used as an indicator of the availability of labile resources [15]. Younger water masses have higher FDOM ratio and lower AOU [15], as a water masses age, microbial activity consumes labile organic compounds and oxygen, leading to higher AOU and lower FDOM ratio.

Differences in the FDOM ratio among water masses were significant (Mann-Whitney test *p* < 0.05) when comparing CDW against NADW, and CDW against WSDW, but not when NADW and WSDW were compared (*p* = 0.49). Furthermore, values of FDOM ratio had positive and significant correlations with viral abundances (Spearman correlation coefficient = 0.48, *p* < 0.05) and virus-to-prokaryote ratios (Spearman correlation coefficient = 0.53, *p* < 0.05). The relative abundance of energy metabolism AMGs was also negatively correlated to the FDOM ratio (Spearman correlation coefficient = −0.32, *p* < 0.05). Analogous to the dichotomy between FL and PA lifestyles, in younger waters masses with higher FDOM ratio, energy is readily available to the microbial hosts through the use of labile organic compounds. Conversely, in older waters masses with lower FDOM ratio, the hosts need to be more efficient at utilizing recalcitrant organic compounds as the main energy source.

We posit a mechanism to explain the higher relative abundances of AMGs involved in the energy metabolism and oxidative phosphorylation observed among older water masses (Figure 6). Previous findings, showed that the genomes of the hosts that thrive within older water masses encode more genes associated with energy metabolism and oxidative phosphorylation, as a mechanism to more efficiently harness energy from the limited supply of organic matter [15]. Thus, the higher relative abundance of AMGs involved in these pathways among the viral genomes could reflect the uptake of metabolic genes from the host genomes. Those viruses that encode such AMGs could have a selective advantage, as they would increase the efficiency of the energy metabolism of their hosts during infection, leading to more viral progeny, and higher relative abundance within the viral community.

**Figure 6:**
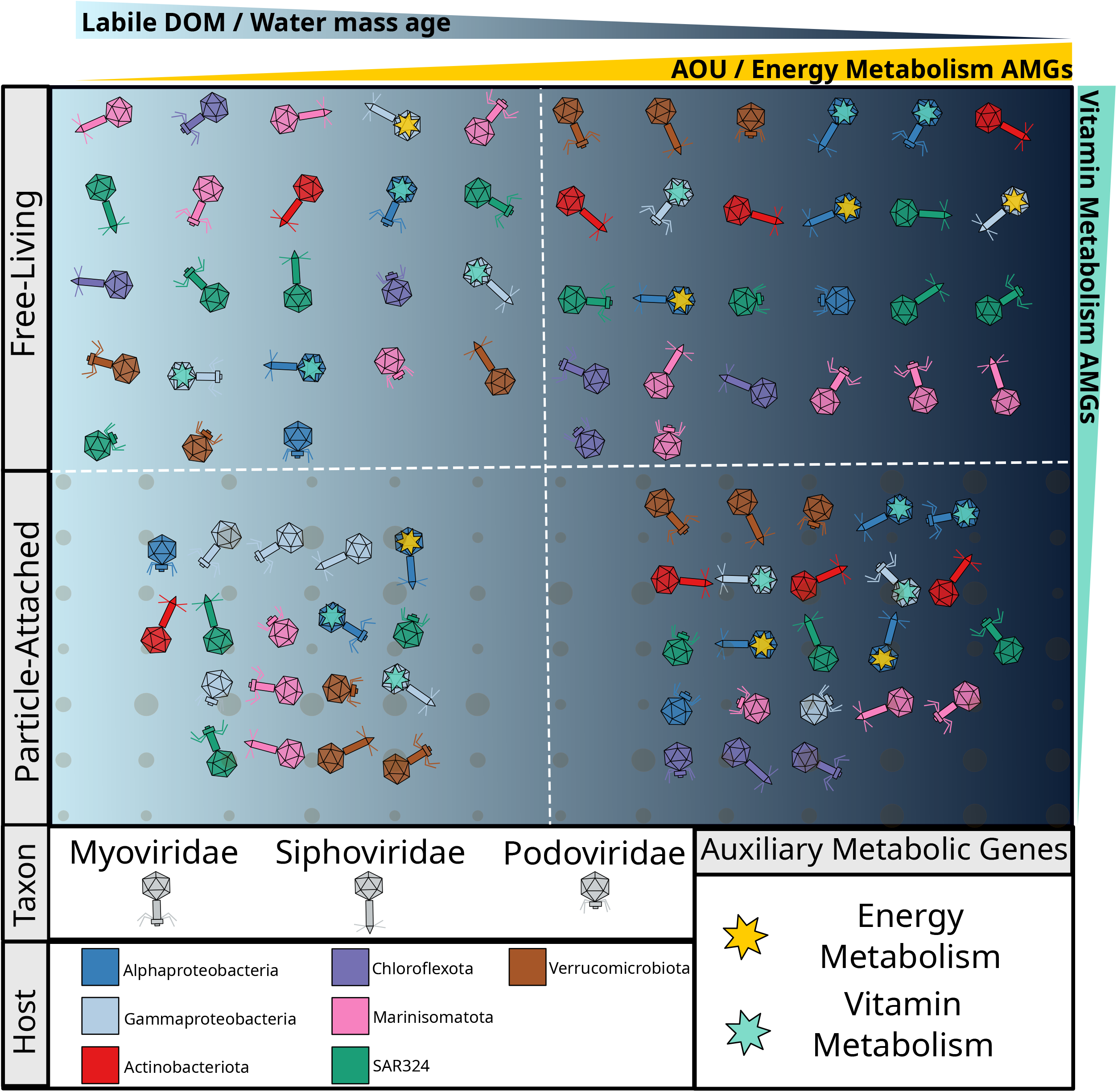
Conceptual model illustrating the changes in viral community between FL and PA and throughout the AOU and FDOM gradients. The most abundant viral families of tailed viruses (Myoviridae, Siphoviridae, and Podoviridae) are depicted according to their morphology. Virus colouring is indicative of predicted host. Auxiliary metabolic genes are depicted by coloured stars inside viral particles.

## Conclusions

Our findings provide insights about deep ocean viruses, one of the least explored biological entities to date. The data corroborates our hypotheses that: 1) Free-living and particle-attached fractions have different viral community compositions; 2) Environmental factors, mainly does related to water mass age, control the abundance, diversity, and AMG repertoire of both free-living and particle-attached viral communities; 3) Particle-attached communities are enriched in viruses that target copiotrophic bacteria, while free-living communities are enriched in viruses that target oligotrophic bacteria; and 4) AMGs from metabolic pathways associated with energy metabolism are enriched in older water masses characterized by lower concentrations of labile organic matter. Our findings lay a solid foundation to the understanding of the factors that structure and influence the functioning of bathypelagic viral communities, by providing a genome, taxonomy, and host-resolved dataset of viruses that includes novel AMG combinations.

## Methods

### Sample collection, environmental parameters

These procedures have previously been described in detail [14]. Briefly, samples were collected as part of the 2010 Malaspina circumnavigation expedition (http://www.expedicionmalaspina.es), which covered both tropical and subtropical regions of the global ocean (Table S1). Measurements of water temperature, salinity, and oxygen concentrations were taken *in situ* [22]. Salinity ranged from 34.64 to 34.91 (median = 34.7, s.d. = 0.09), temperature ranged from 0.58 to 2.25 °C (median = 1.5, s.d. = 0.48).

Apparent oxygen utilization (AOU) is the parameter used to estimate water age. It is calculated as the difference between oxygen solubility in a water mass and its measured oxygen concentration, integrating all respiratory processes since the last contact of the water mass with the atmosphere. Thus, AOU was measured as the difference between oxygen solubility in water samples and its measured oxygen concentration. Oxygen solubility is determined by pressure, water temperature, and salinity. Older water masses have lower oxygen concentrations and higher AOU and vice-versa. AOU ranged from 97 to 231 μmol/kg (median = 163, s.d. =39). There were significant (*p* < 0.01, Pearson product-moment correlation) associations between salinity, temperature and AOU.

Water samples were retrieved to characterize the Fluorescent Dissolved Organic Matter (FDOM) content of the water masses from which the metagenomes were obtained [28,39]. The FDOM composition is characterized by a pair of recalcitrant humic-like compounds (C1 and C2), and a pair of comparatively more labile compounds (C3 and C4), which are commonly attributed to the amino acids tryptophan and tyrosine respectively [39]. The FDOM ratio was calculated as (C3 + C4) /(C1 + C2). This variable is a proxy for the proportion of labile compounds in the dissolved organic matter pool [15].

### Metagenome and viral genome analysis

For each sample, 120 liters of seawater were sequentially filtered through 200-20 μm meshes to enrich the samples for bacteria, archaea and viruses. Next, size fractioning was used to separate the free-living (0.2–0.8 μm) and particle-attached (0.8–20 μm) fractions. Filtered samples were flash frozen in liquid nitrogen until DNA extraction which was performed through the phenolchloroform method. DNA was sequenced at the DOE’s Joint Genome Institute (JGI) in an Illumina HiSeq2000 platform. Post-QC reads from metagenomes were co-assembled using Megahit v1.2.8 [40]. Sequences shorter than 2 Kbp as well as redundant sequences were removed with CD-HIT [41].

All of the following analyses were performed with default parameters unless otherwise stated. All the assembled scaffolds derived from the co-assembly of metagenomes were processed through VIBRANT v1.2.1 [30] to identify viral sequences. The quality of the obtained viral genomes and genome fragments was assessed with CheckV v0.7.0 [31]. Taxonomic classification of viral sequences was achieved through VPF-Class version dd88a54 [32]. Viral host predictions were performed using PHIST version ed2a1e6 [33], the previously described set of metagenome assembled genomes from the Malaspina dataset [14], were used as putative hosts. For the PHIST analysis, only predictions with a maximum e-value of 2.384e-14 were considered, which yields approximately 85% class level prediction accuracy.

Coding DNA Sequences (CDS) in viral genomes were identified with Prodigal v2.6.3 [42]. Viral CDS were queried against a large database containing protein sequences derived from approximately 4.6 million viral genomes and genome fragments derived from the latest release of IMG/VR [43] and complemented with sequences from multiple ecosystems. Protein searches were performed using MMSeqs2 [44]. Next, the pairwise similarity scores were calculated between query and reference genomes based on the number of matched CDS, percentage of matched CDS and average amino acid identity (AAI). To exclude spurious similarities the following thresholds were applied: a minimum of 3 CDS matches, covering at least 30% of all CDS in the query scaffold, and yielding a minimum of 30% AAI. CDS were queried against three databases for functional taxonomic annotation: 1) UniRef100 using DIAMOND version 2.0.7 [45], 2) KOFam using Hmmscan version 3.3 [46], 3) Pfam using Hmmscan as well. For all searches only hits the displayed a bitscore ≥ 50 and e-value ≤ 10^-5^ were considered for subsequent analysis.

### Viral community composition analysis

Post-QC reads from the metagenomes were queried against the database of obtained viral genomes using Bowtie v2.3.4.1 [47] in sensitive local mode. Genome abundances were calculated as Reads Per Kilobase per Million total sequences (RPKM). Viral genome abundances were grouped by viral families or predicted host taxon as the sum of RPKM values of all the viruses in each group [2,48]. Similarly, abundances of KEGG KOs were calculated as the sum of RPKM values of all genomes encoding a given KO. Finally, to calculate the abundances of KEGG pathways and metabolisms we summed the abundances of all genomes encoding a given KO associated with a metabolism/pathway (Table S5). To avoid overestimating abundances for KOs that belong to multiple metabolisms/pathways, the abundance of each KO was divided by the number of metabolisms/pathways the KO was assigned before calculating sums (Tables S6 and S7).

### Statistical analysis

Sample community composition analyses were performed based on calculated RPKM values of viral scaffolds. Non-metric Multidimensional Scaling (NMDS) was performed by first calculating Bray-Curtis distances between communities within the Vegan package [49]. Permutational multivariate analysis of variance (PERMANOVA) was conducted with 1,000 permutations and Bray-Curtis distances. Differences in variable relative abundances (i.e., taxa, predicted hosts, or functions) between free-living and particle-attached fractions were evaluated with the Mann-Whitney test. Associations between environmental and metagenome variables were assessed by calculating either Pearson or Spearman correlation scores. For both correlation and Mann-Whitney tests associations with a *p-value* ≤ 0.05 were considered significant. Multiple testing correction was performed through the Bonferroni method. All analyses were carried out in R v4.0.0 [50].

## Acknowledgments

We thank the R/V Hespérides captain and crew, the chief scientists in Malaspina expedition legs, and all project participants for their help in making this project possible. This work was funded by the Spanish Ministry of Economy and Competitiveness (MINECO) through the Consolider-Ingenio program (Malaspina 2010 Expedition, ref. CSD2008-00077). The sequencing of 58 bathypelagic metagenomes was done by the U.S. Department of Energy Joint Genome Institute, supported by the Office of Science of the U.S. Department of Energy under Contract No. DE-AC02 05CH11231 to SGA (CSP 612 “Microbial metagenomics and transcriptomics from a global deep-ocean expedition”). Additional funding was provided by the project MAGGY (CTM2017-87736-R) to S.G.A. from the Spanish Ministry of Economy and Competitiveness, Grup de Recerca 2017SGR/1568 from Generalitat de Catalunya, and King Abdullah University of Science and Technology (KAUST) under contract OSR #3362). The ICM researchers have had the institutional support of the “Severo Ochoa Centre of Excellence” accreditation (CEX2019-000928-S). High-Performance computing analyses were run at the Marine Bioinformatics Service (MARBITS, https://marbits.icm.csic.es) of the Institut de Ciències del Mar (ICM-CSIC). FHC was supported by a Juan de la Cierva - Incorporación fellowship (Grant IJC2019-039859-I).

## Data availability statement

Identified viral scaffolds were deposited in ENA under project ID PRJEB40454

## Code availability statement

The code available at https://github.com/felipehcoutinho/virathon

## Author contributions

SGA, JMG, DV, and CD coordinated the sampling expedition. FHC, CS, and MS designed the data analysis protocol. FHC wrote the code used to analyse the metadata and genomic data. FHC, PS, and MS analysed the results. FHC wrote the manuscript. All authors contributed to reviewing the manuscript.

## Ethics Approval and Consent to Participate

Not Applicable.

## Consent for publication

Not Applicable.

## Supplementary Material

**Figure S1:** Relative abundance patterns of viral scaffolds encoding DHFR genes. A) Stacked bar plots depicting the RPKM abundances (y-axis) of viral scaffolds across samples (x-axis), separated by free-living and particle-attached samples (panels). Sampling stations are sorted from left to right by increasing oxygen concentrations. B) Box plots depicting the differences in DHFR encoding scaffold abundances between free-living and particle-attached fractions. Boxes depict the median, the first and third quartiles. Whiskers extend to 1.5 of the interquartile ranges. Outliers are represented as dots above or below whiskers. The *p*-values of each comparison obtained with the Mann-Whitney test are depicted above bars.

**Table S1:** Metadata of the collected samples and metagenomes.

**Table S2:** Metadata of the obtained viral genomes and genome fragments.

**Table S3:** Functional and taxonomic annotation of the CDS encoded in the viral genomes and genome fragments.

**Table S4:** Relative abundances of viral scaffolds across 58 metagenomes calculated as RPKM.

**Table S5:** Relative abundances of viral KEGG KOs across 58 metagenomes calculated based on RPKM of individual viral scaffolds.

**Table S6:** Relative abundances of viral KEGG pathways across 58 metagenomes calculated based on RPKM of individual viral scaffolds.

**Table S7:** Relative abundances of viral KEGG metabolism across 58 metagenomes calculated based on RPKM of individual viral scaffolds.

**Table S8:**Relative abundances of viruses grouped according to family level taxonomic affiliation based on RPKM of individual viral scaffolds.

**Table S9:** Relative abundances of viruses grouped according to predicted host phylum (or class for Proteobacteria) taxonomic affiliation based on RPKM of individual viral scaffolds.

