## Supplementary figures and images for "Water mass age structures the auxiliary metabolic gene content of free-living and particle-attached deep ocean viral communities"

### Figure S1

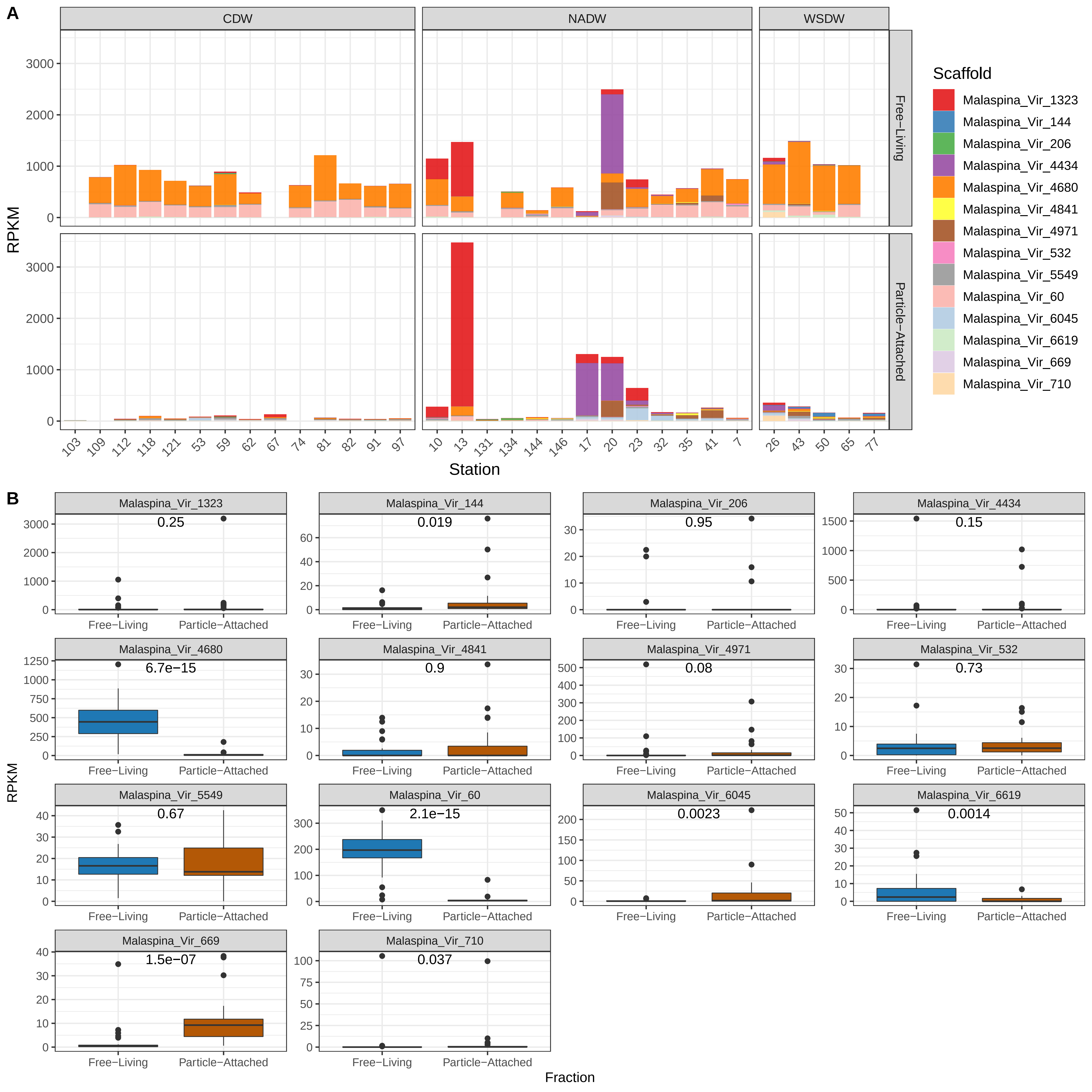
